## Supplementary Figures for "Mitochondrial ATP synthesis is essential for efficient gametogenesis in *Plasmodium falciparum*"

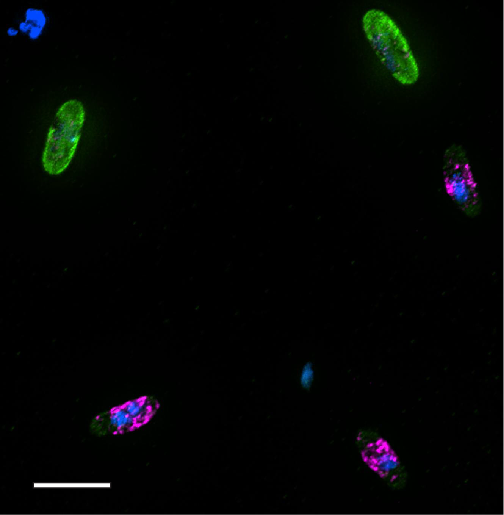

**Supplementary Figure 1** – Male and female Stage V gametocytes stained with the putative male gametocyte marker anti-LDH2 (green), the established female marker anti-PfG377 (magenta) and DAPI (blue). Scale bar = 12 µm.

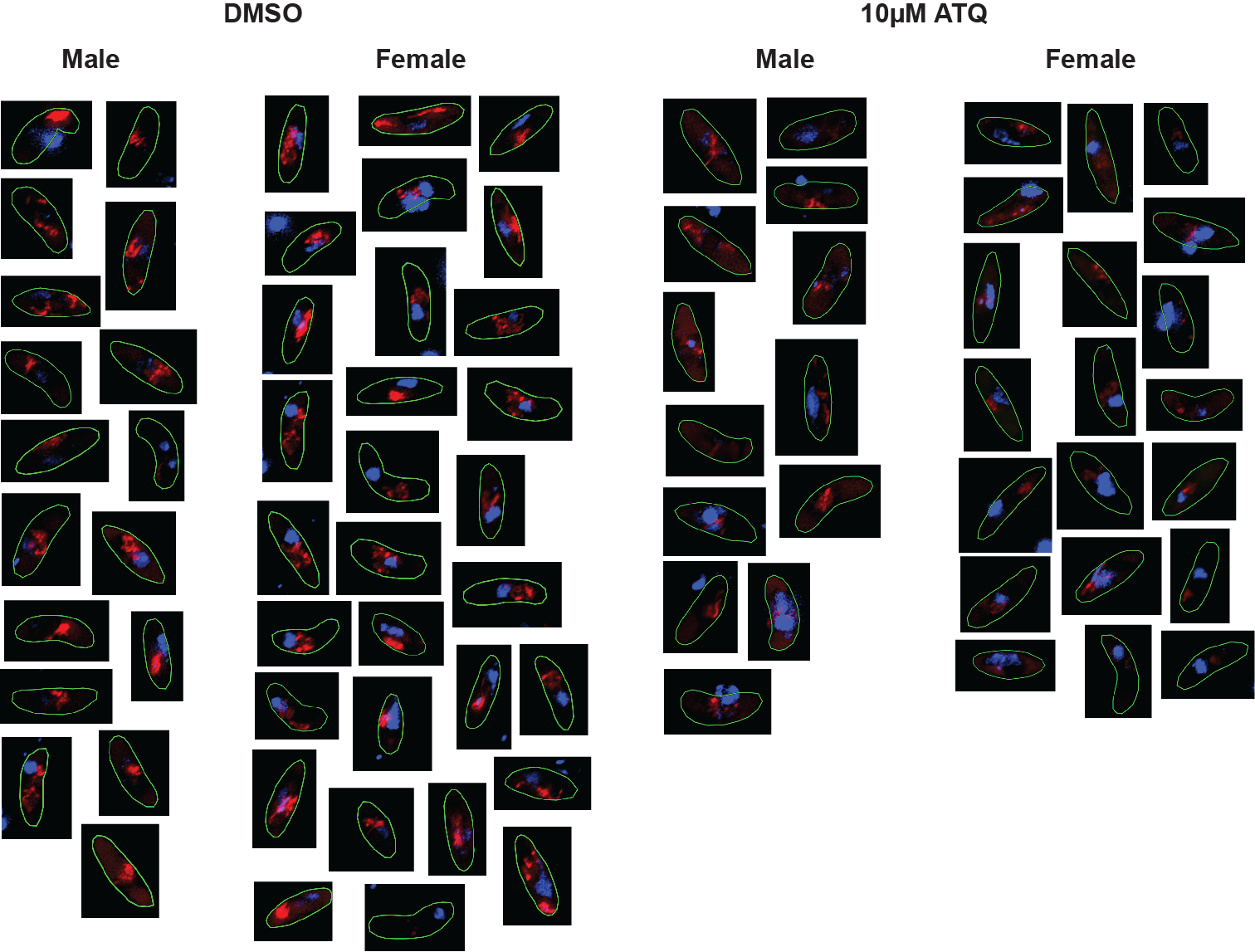
 **Supplementary Figure 2** – A montage of individual male and female gametocytes from the analysis in Figure 2 and Figure 3. Shown here are the outline of the cell (green), MitoTracker staining (red) and DAPI staining (blue). Mitochondrial morphology varies substantially between cells, with no clear pattern distinguishing male and female gametocytes. Gametocytes treated with 10µM atovaquone (ATQ) show reduced MitoTracker incorporation.

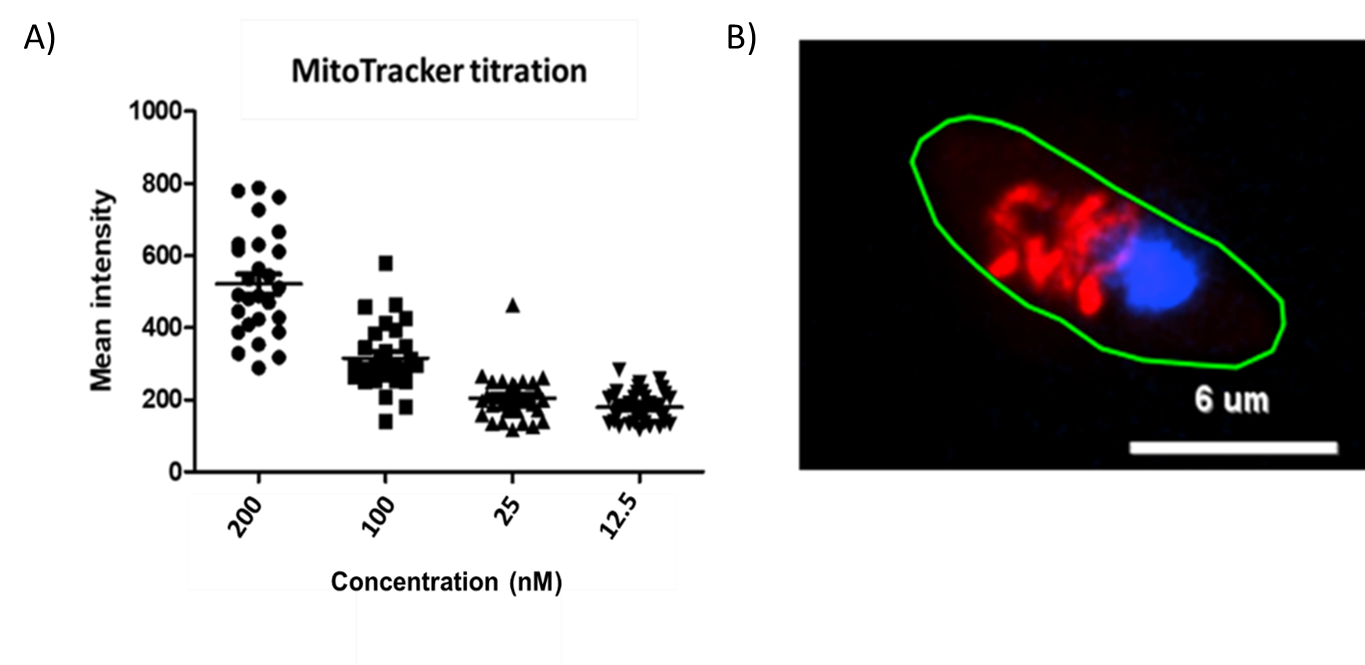

**Supplementary Figure 3** – Live gametocytes were treated with decreasing concentrations of Mitotracker for 25 minutes before being washed and fixed. Intensity of mitotracker staining was then calculated for individual cells by fluorescence imaging and quantitative image analysis. A detectable signal was observed in gametocyte mitochondria treated with as little as 12.5nM Mitotracker.

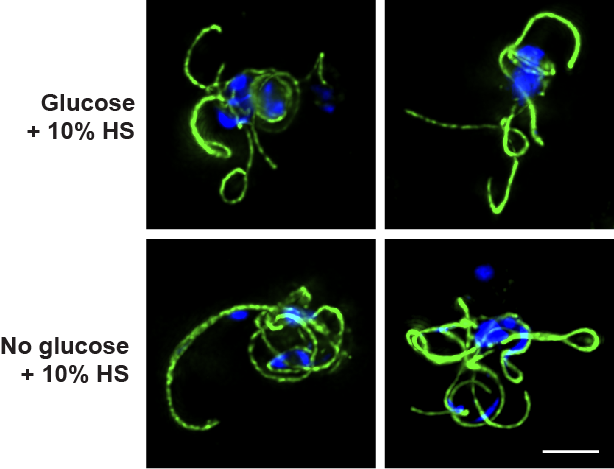

**Supplementary Figure 4 – Short term incubation in glucose-free RPMI has no effect on male gametogenesis.** Gametocytes were incubated for 1 h with culture medium containing RPMI with or without glucose, plus 10% human serum (HS). They were then triggered in condition-matched ookinete medium, fixed 20 min later, and stained with anti-aplha tubulin II (green) and DAPI (blue). We conclude that 10% human serum contains sufficient endogenous glucose to fuel exflagellation. Scale bar = 5 µM
